# Sharing brain mapping statistical results with the neuroimaging data model

**DOI:** 10.1101/041798

**Authors:** Camille Maumet, Tibor Auer, Alexander Bowring, Gang Chen, Samir Das, Guillaume Flandin, Satrajit Ghosh, Tristan Glatard, Krzysztof J. Gorgolewski, Karl G. Helmer, Mark Jenkinson, David B. Keator, B. Nolan Nichols, Jean-Baptiste Poline, Richard Reynolds, Vanessa Sochat, Jessica Turner, Thomas E. Nichols

## Abstract

Only a tiny fraction of the data and metadata produced by an fMRI study is finally conveyed to the community. This lack of transparency not only hinders the reproducibility of neuroimaging results but also impairs future meta-analyses. In this work we introduce NIDM-Results, a format specification providing a machine-readable description of neuroimaging statistical results along with key image data summarising the experiment. NIDM-Results provides a unified representation of mass univariate analyses including a level of detail consistent with available best practices. This standardized representation allows authors to relay methods and results in a platform-independent regularized format that is not tied to a particular neuroimaging software package. Tools are available to export NIDM-Result graphs and associated files from the widely used SPM and FSL software packages, and the NeuroVault repository can import NIDM-Results archives. The specification is publically available at: http://nidm.nidash.org/specs/nidm-results.html.

## Introduction

A neuroimaging technique like functional Magnetic Resonance Imaging (fMRI) generates hundreds of gigabytes of data, yet only a tiny fraction of that information is finally conveyed to the community. In a typical paper, the entire results report consists of 1) a list of significant local maxima, i.e. locations in the brain defined in a standard atlas space inferred to be distinct from noise, and 2) a graphical representation of the activations as an image figure.

This practice is unsatisfactory for three reasons. First, because it represents a massive loss of information from the raw and even the derived data used to draw the conclusion of the study. For example, a meta-analysis (in settings other than neuroimaging) combines estimates of effects of interest and their uncertainty across studies. In brain imaging, the locations of local maxima have no measures of uncertainty reported. While neuroimaging meta-analysis methods for coordinate data exist ^1–3^ they are a poor approximation to the meta-analysis that would be obtained if the image data were available ^4^. Even though there are emerging infrastructures to support sharing of neuroimaging data (e.g. NeuroVault RRID:SCR_003806 ^5,6^), these are still rarely utilised ^7^.

Second, despite the availability of guidelines ^8–10^, ambiguous or incomplete methodological reporting in papers is still commonplace ^11^ hindering the robustness and reproducibility of scientific results ^11,12^.

Finally, key methodological details of the study are described in free-form text in a paper and not available in machine-readable form, making these metadata essentially unsearchable. Databases have been built to provide metadata associated with published papers, either manually curated (e.g. BrainMap ^13,14^) or automatically-populated using text-mining algorithms (e.g. NeuroSynth ^15,16^), but, ideally, these metadata should be made available by the authors themselves at the time of the publication, together with the data. Additionally, searchable metadata, could help identify potential confounding factors that are currently being overlooked (e.g. how different smoothing kernels impact the meta-analysis, or the influence of different processing strategies on the outcome of the analysis).

In order to make neuroimaging results available in a machine-readable form a number of key technical issues have to be addressed. First, the scope of the metadata to be shared must be defined. The space of possible metadata to report is extremely large encompassing experimental design, acquisition, pre-processing, statistical analysis, etc. The optimal set of metadata is highly dependant on the application of interest and possible applications of shared data are broad. For example, in a meta-analysis, the contrast standard error map is required, while a comparison across neuroimaging processing pipelines would require a complete description of the analysis pipeline including software-specific parameterization.

Another technical issue is the need to define a common representation across neuroimaging software packages. While the three main neuroimaging software packages, SPM (RRID:SCR_007037) ^17,18^, FSL (RRID:SCR_002823) ^19,20^ and AFNI (RRID:SCR_005927) ^21,22^, all implement similar analyses, they often use different terms to refer to the same concept. For example, FSL’s parameter estimate maps (e.g. pe1.nii.gz) are the equivalent of SPM’s beta maps (e.g. beta_0001.nii). They can also use the same term when referring to different concepts. For example, SPM and FSL both use a global scaling of the data to get “percent BOLD signal change”, but due to differences in how the brain mask and mean signal are computed, the data are scaled quite differently ^23^ and are not comparable. In order to fully describe an analysis, the sharing of software-specific batch scripts (e.g. SPM matlabbatch files, FSL fsf files, or history stored in AFNI brick headers) would be a simple solution to provide all the parameters from an analysis, but the ability to compare and query across software would still be lacking. Pipeline systems like NiPype ^24^, LONI Pipeline ^25^ and CBRAIN ^26^ do explicitly model analysis steps, but a large volume of research is still conducted directly with tools not embedded in pipelines. Ideally, one should be able to identify all studies corresponding to a set of criteria of interest regardless of the software used. This will only be possible if information about results across software can be represented using common data elements and structures.

A machine-readable representation of neuroimaging data and results, using a common descriptive standard across neuroimaging software packages, would address these issues of comparability and transparency.

A previous effort in this direction was the XML-based Clinical and Experimental Data Exchange (XCEDE) schema ^27^, developed in the context of the Biomedical Informatics Research Network (BIRN) ^28,29^. XCEDE modelled information about both the experimental design and results (peaks and clusters) in neuroimaging studies. This XML schema was defined to be independent from any particular neuroimaging analysis software and was made openly available ^30^. XCEDE has been used by multiple sites across the United States and the United Kingdom in the context of the fBIRN project and is still in use by the Human Imaging Database ^28,31^. An implementation was provided for SPM ^32,33^ as well as a set of tools ^34^. However, the XCEDE model was not implemented by other imaging software, supported limited provenance information, and did not offer the ability to jointly share image data summarising the experiment.

Beyond neuroimaging, encoding of provenance, i.e. keeping track of the processes that were applied to the data, encompassing a description of the tools, data flow and workflow parameterization, is a topic of growing interest in science in general. A number of solutions have been proposed in order to support better documentation of research studies. Among them, the PROV data model ^35^ is a W3C specification to describe provenance on the web. PROV is defined in a generic fashion that is not tied to a domain in particular (cf. ^36^ for examples of implementations).

The NeuroImaging Data Model (NIDM) ^37,38^ was created to expand upon the initial development of XCEDE, introducing a domain-specific extension of PROV using semantic web technologies and the Resource Description Framework (RDF). The goal of NIDM is to provide a complete description of provenance for neuroimaging studies, from raw data to the final results including all the steps in-between. The core motivation of NIDM is to support data sharing and data reuse in neuroimaging by providing rich machine-readable metadata. Since its first developments in 2011, NIDM has been an ongoing effort and is currently comprised of three complementary projects: NIDM-Experiment, NIDM-Workflows and NIDM-Results. NIDM-Experiment targets the representation of raw data generated by the scanner and information on the participants. NIDM-Workflows focuses on the description of data analysis parameterization, including detailed software-specific variations. NIDM-Results, presented here, deals with the representation of mass-univariate neuroimaging results using a common descriptive standard across neuroimaging software packages.

A motivating use case for NIDM-Results was neuroimaging meta-analysis, but the format also produces a detailed machine-readable report of many facets of an analysis. The implementation of NIDM-Results within SPM and FSL, two of the main neuroimaging software packages, provides an automated solution to share maps generated by neuroimaging studies along with their metadata. While NIDM-Results focuses on mass-univariate studies and is mostly targeted at fMRI, the standard is also suitable for anatomical MRI (with Voxel-Based Morphometry), and Positron Emission Tomography (PET). It was developed under the auspices of the International Neuroinformatics Coordinating Facility (INCF) Neuroimaging data sharing Task Force (NIDASH) which comprises a core group of experts representing more than ten labs involved in various facets of neuroimaging (including statistical analysis, informatics, software development, ontologies). It also involved close collaboration with the main neuroimaging software developers. The format is natively implemented in SPM and a NIDM-Results exporter is available for FSL and will be integrated in a future version of FSL. Both NeuroVault and CBRAIN support export to NIDM-Results and NeuroVault additionally can import NIDM-Results archives.

## Results

### Model

#### Definitions

The definitions provided below are used throughout the manuscript:

- **NIDM-Results graph**: A particular instance of a representation of data and metadata complying with the NIDM-Results specification.
- **NIDM-Results serialization**: A text file rendering of a NIDM-Results graph.
- **NIDM-Results pack**: A compressed file containing a NIDM-Results serialization and some or all of the referenced image data files.

#### Overview

The NIDM-Results standard is defined by a W3-style specification, publicly available at http://nidm.nidash.org/specs/nidm-results.html and by an ontology (owl) file available at http://bioportal.bioontology.org/ontologies/NIDM-RESULTS. It is comprised of a controlled vocabulary, as well as instructions of how to use PROV to represent mass-univariate neuroimaging results. The model provides terms to describe key elements of neuroimaging methods using a common framework across neuroimaging software packages. For example, as illustrated in Fig. 1, error models are described in terms of assumed variance (homoscedastic, heteroscedastic) and assumed covariance structure (independent, spatially correlated, etc.) and how these structures vary in space (defined independently at each voxel, globally throughout the brain or spatially regularised).

**Fig. 1.**
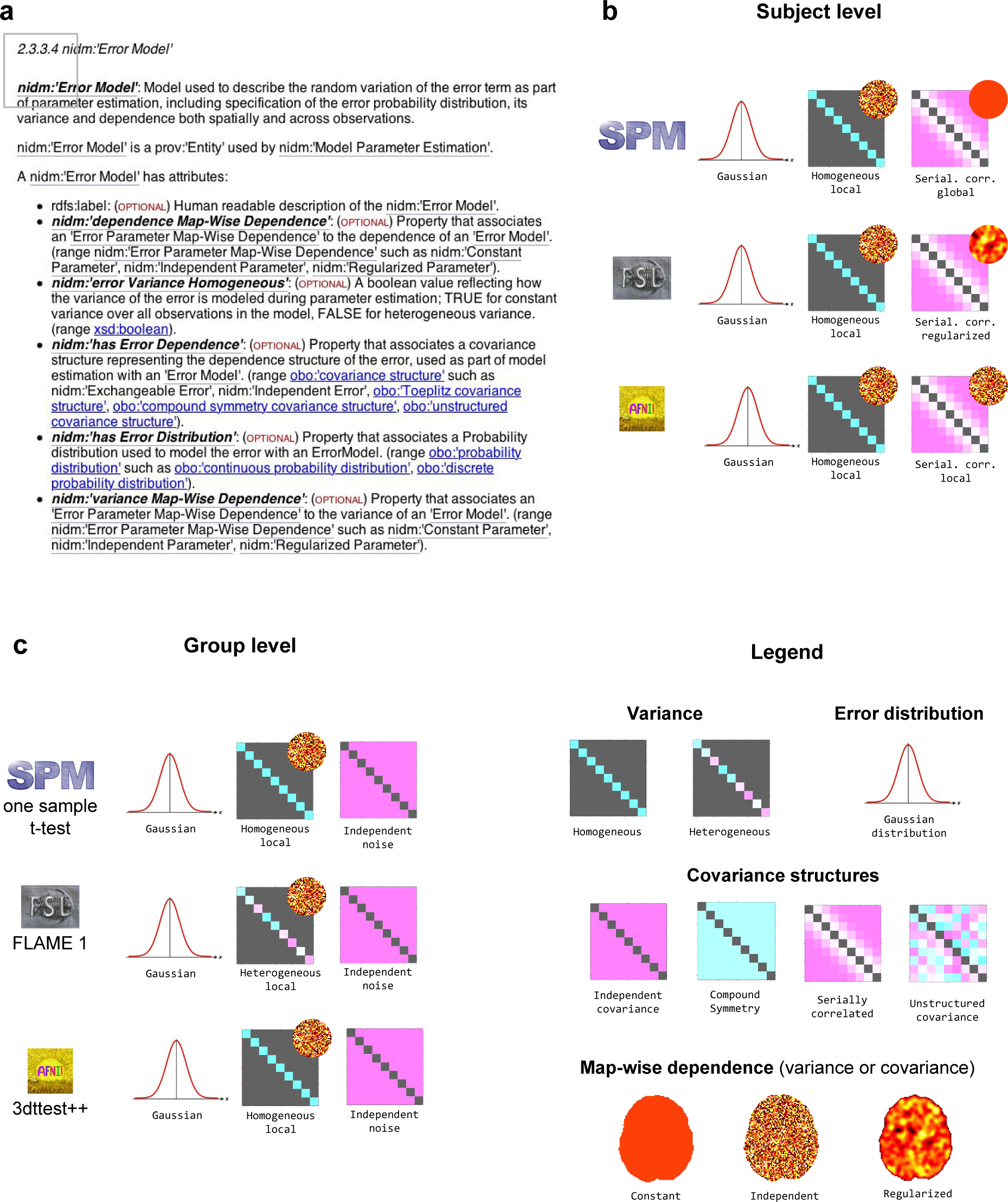
Description of the error models with NIDM-Results. Excerpt of the NIDM-Results 1.3.0 specification describing a nidm:’Error Model’ and its attributes (a). Examples of model implementations for subject-level (b) and group-level (c) analyses for SPM, FSL and AFNI.

The current version, NIDM-Results 1.3.0, defines 214 terms (140 classes and 74 attributes) of which 45 terms are re-used from external vocabularies and ontologies. All terms are defined as specialisations of the PROV terms. Three namespaces are defined: http://purl.org/nidash/nidm#(“nidm:”), http://purl.org/nidash/spm#(“spm:”) and http://purl.org/nidash/fsl#(“fsl:”). Anything that could be represented across software or that is a generic concept is defined in the “nidm:” namespace. Software-specific namespaces: (“spm:”, “fsl:”) are reserved for the description of functionality unique to one software (e.g. global null inference for conjunction testing in SPM).

Fig. 2 provides an overview of NIDM-Results. In the description below, terms in single quote correspond to elements defined by the model, identifiers for those terms are provided in Table 1.

**Fig. 2.**
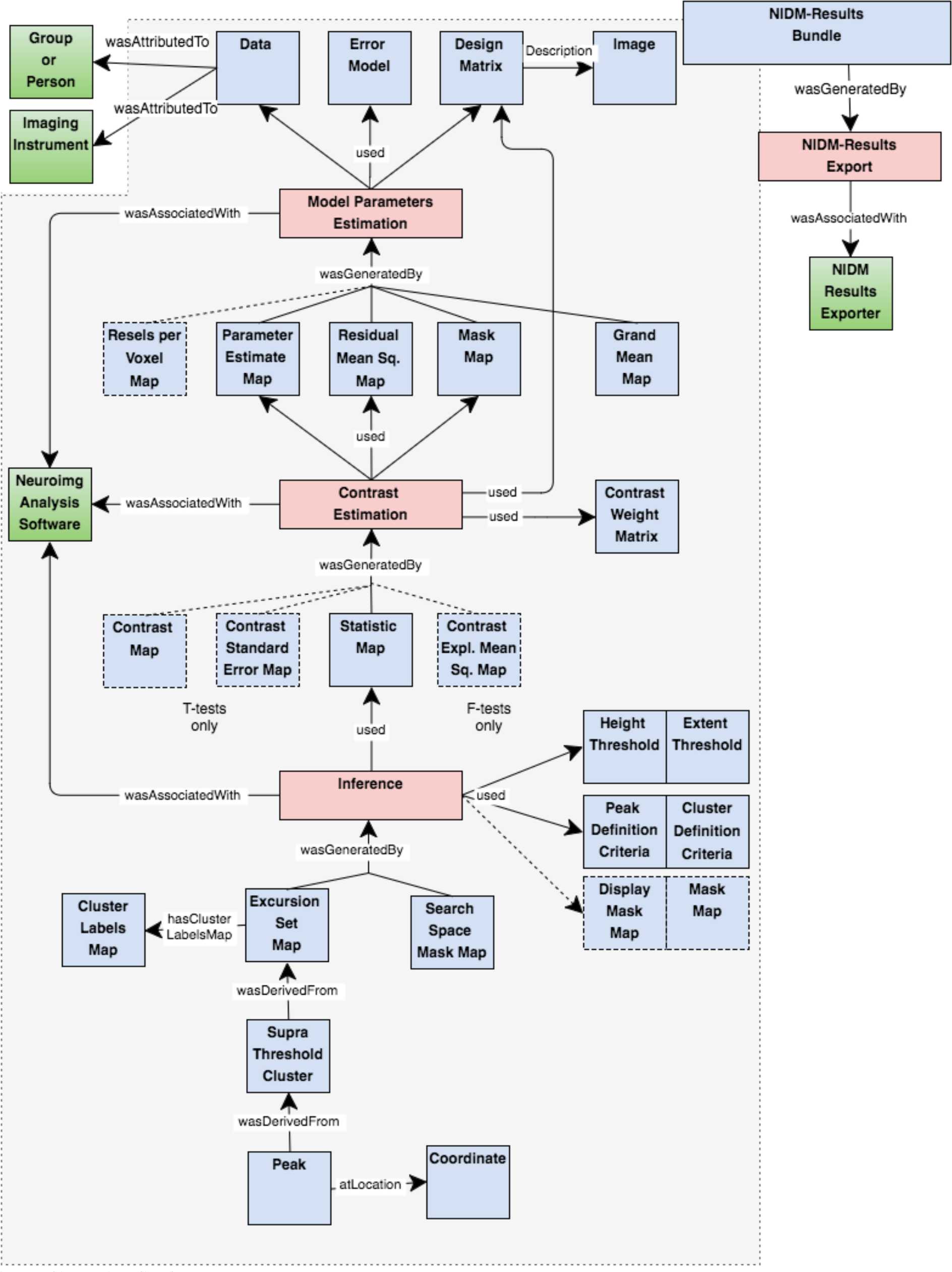
NIDM-Results objects. Color-coding indicates the type as defined in PROV (blue: Entity, red: Activity, green: Agent).

**Table 1.** PROV type, label and identifier of the NIDM-Results terms mentioned in single quotes in this manuscript.

| PROV type | Term | Qualified name |
| --- | --- | --- |
| Entity | NIDM-Results bundle | nidm:NIDM_0000027 |
|  | Bundle | prov:Bundle |
|  | Design Matrix | nidm:NIDM_0000019 |
|  | Error Model | nidm:NIDM_0000023 |
|  | Data | nidm:NIDM_0000169 |
|  | Parameter Estimate Map(s) | nidm:NIDM_0000061 |
|  | Mask Map | nidm:NIDM_0000054 |
|  | Residual Mean Squares Map | nidm:NIDM_0000066 |
|  | Resels Per Voxel Map | nidm:NIDM_0000144 |
|  | Grand Mean Map | nidm:NIDM_0000033 |
|  | contrast weight matrix | obo:STATO_0000323 |
|  | Statistic Map | nidm:NIDM_0000076 |
|  | Contrast Map | nidm:NIDM_0000002 |
|  | Contrast Standard Error Map | nidm:NIDM_0000013 |
|  | Contrast Explained Mean Square Map | nidm:NIDM_0000163 |
|  | Excursion Set Map | nidm:NIDM_0000025 |
|  | Height Threshold | nidm:NIDM_0000034 |
|  | Extent Threshold | nidm:NIDM_0000026 |
|  | Peak Definition Criteria | nidm:NIDM_0000063 |
|  | Cluster Definition Criteria | nidm:NIDM_0000007 |
|  | Display Mask Map | nidm:NIDM_0000020 |
|  | Search Space Mask Map | nidm:NIDM_0000068 |
|  | Supra-Threshold Cluster(s) | nidm:NIDM_0000070 |
|  | Peak(s) | nidm:NIDM_0000062 |
| Activity | Model Parameter Estimation | nidm:NIDM_0000056 |
|  | Contrast Estimation | nidm:NIDM_0000001 |
|  | Inference | nidm:NIDM_0000049 |
|  | Conjunction Inference | nidm:NIDM_0000011 |
|  | NIDM-Results Export | nidm:NIDM_0000166 |
| Agent | Neuroimaging Analysis Software | nidm:NIDM_0000164 |
|  | Person | prov:Person |
|  | study group population | obo:STATO_0000193 |
|  | Imaging Instrument | nif:birnlex_2094 |
|  | NIDM-Results Exporter | nidm:NIDM_0000165 |
|  | nidmfsl | nidm:NIDM_0000167 |
|  | spm_results_nidm | nidm:NIDM_0000168 |

The main entity is a *‘NIDM-Results bundle’*, a specialisation of a ‘Bundle’ as defined in PROV, i.e. an entity gathering a set of entities, activities and agents. A *‘NIDM-Results bundle’* contains a description of the mass univariate results provenance and is typically made up of:

- 3 activities representing the main steps of statistical hypothesis testing: ‘*model parameter estimation’*, ‘*contrast estimation’* and ‘*inference’*.
- 26 types of entities (of which 6 are optional) representing inputs and outputs of the activities;
- 3 agents representing the ‘*neuroimaging analysis software’*, the *‘person’* or *‘study group population’* who participated in the study and the type of *‘imaging instrument’* used.

The statistical model is described in the ‘*design matrix’* and ‘*error model’* entities that are both used by the ‘*model parameter estimation’* activity. The *‘data’* entity describes the scaling applied to the data before model fitting (especially relevant for first-level fMRI experiments) and links to the participants (as a ‘person’ or a group) and the ‘imaging instrument’ used to acquire the data (e.g. a magnetic resonance imaging scanner or an electroencephalography machine). A set of ‘*parameter estimate maps’* is generated by the ‘*model parameter estimation’* activity along with the analysis ‘*mask map’*, a ‘*residual mean squares map’* and a ‘*grand mean map’* that can be used to check the performance of the data scaling. Optionally, a *‘resels per voxel map’* can also be generated to record local variations in noise smoothness.

The ‘*contrast estimation’* activity uses a subset of the ‘*parameter estimate maps’*, the ‘*residual mean squares map’* and the analysis ‘*mask map’* and combine them according to a *‘contrast weight matrix’* to generate a ‘*statistic map’*. For T-tests, a ‘*contrast map’* along with its ‘contrast *standard error map’* are also generated while for F-tests a ‘*contrast explained mean square map’* (i.e. the numerator of an F-statistic) is provided.

Finally, the ‘*inference’*activity uses a ‘*statistic map’* and generates an ‘*excursion set map’* given a ‘*height threshold’* and an ‘*extent threshold’*. The ‘*peak definition criteria’* and ‘*cluster definition criteria’* entities, used by ‘*inference’*, provide the connectivity criterion and minimal distance between peaks (e.g. default is set to 8 mm for SPM and 0 mm for FSL). The ‘*inference’* activity can be replaced by a ‘*conjunction inference’* which uses more than one statistic map. An optional ‘*display mask map’* entity can be used to represent contrast masking, i.e. to restrict the display without affecting the correction for multiple comparisons. The ‘*inference’* activity also generates the ‘*search space mask map’* that represents the search region in which the inference was performed (i.e. the intersection of all input mask maps, except for the display mask map). A set of ‘*supra-threshold clusters’* is derived from the ‘*excursion set map’*and a set of ‘*peaks’* is derived from each cluster. Those are the clusters and peaks that are typically reported in the results of a neuroimaging study.

A *‘neuroimaging analysis software’* agent represents the software package used to compute the analysis. This agent is associated with all activities within the bundle.

Provenance of the *‘NIDM-Results bundle’* is also recorded: the bundle was generated by a *‘NIDM-Results Export’* activity which was performed by a *‘NIDM-Results Exporter’* software agent corresponding to the software used to create the NIDM-Results document (e.g. FSL’s Python scripts, named *‘nidmfsl’* or SPM’s exporter named *‘spm_results_nidm’*). The bundle is associated with a version number corresponding to the version of NIDM-Results model in use.

Each activity, entity and agent has a number of predefined attributes. For instance, the list of attributes of an ‘*error model’* entity is provided in Fig. 1.

#### Updates

Each release of NIDM-Results is associated with a version number. Comments on the current version as well as suggestions of extension can be provided on the GitHub nidm repository: https://github.com/incfnidash/nidm. Each extension or proposition of update will be reviewed and discussed with the members of the INCF NIDASH task force.

### Implementation

SPM12 natively supports export of its results into a NIDM-Results pack, either by the use of a contextual menu in the results table or non-interactively via the batch interface as illustrated in Fig. 3. Export of FEAT results from FSL into a NIDM-results pack can be performed using the Python module nidmfsl (https://pypi.python.org/pypi/nidmfsl), as also illustrated in Fig. 3. nidmfsl was integrated in NeuroVault and as a plugin ^39^ of the CBRAIN web platform for high-performance computing (RRID:SCR_005513) ^26^. As a result, any FSL FEAT analysis uploaded to NeuroVault or performed in CBRAIN can be exported as a NIDM-Results pack. NeuroVault also accepts NIDM-Results packs as a mean to upload new data to a collection. The nidmresults Python library (https://pypi.python.org/pypi/nidmresults) and the provenance MATLAB toolbox (http://www.artefact.tk/software/matlab/provenance/) provide higher-level functions to interact with NIDM-Results packs.

**Fig. 3.**
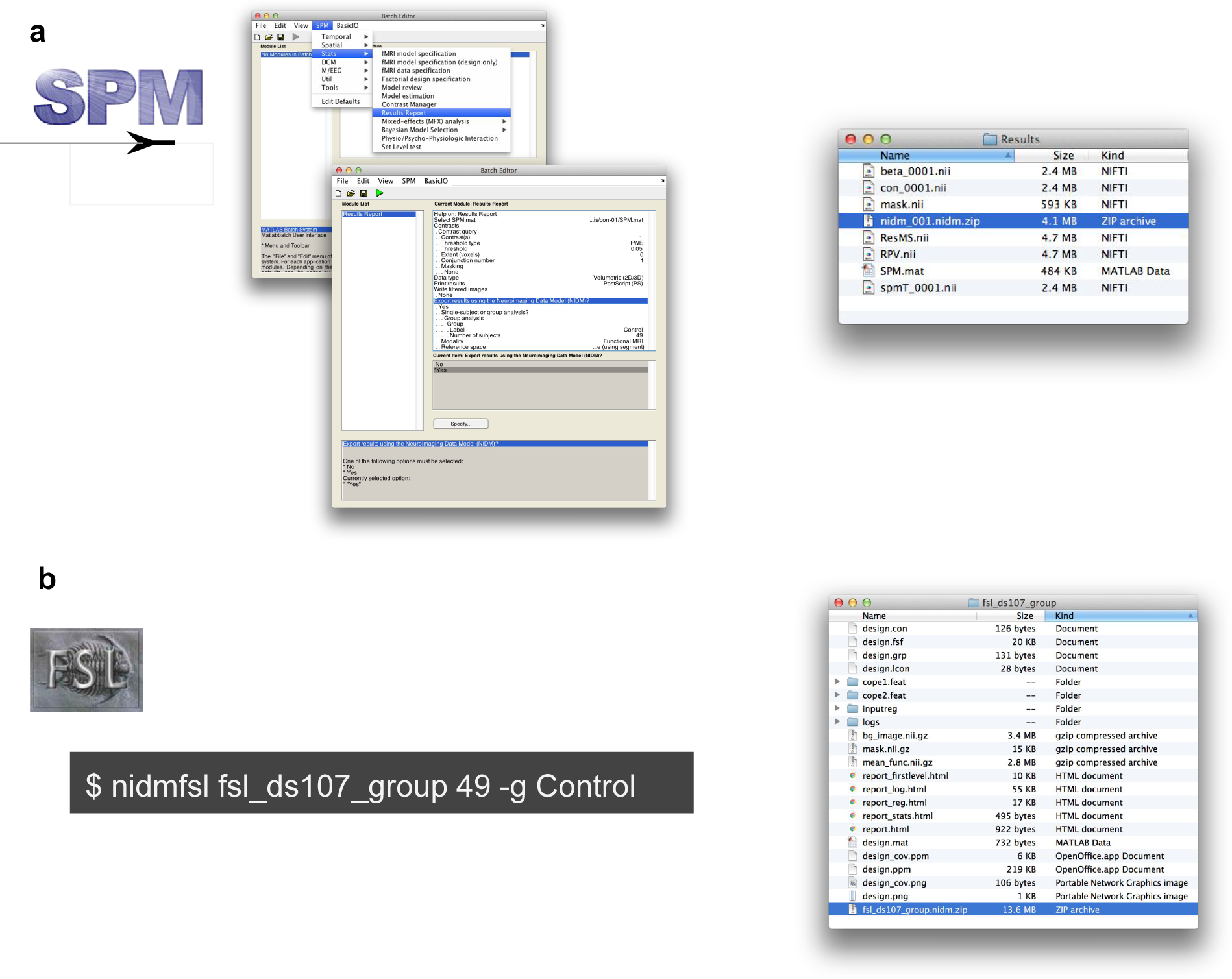
NIDM-Results export in SPM12 (a) and FSL v5.0 (b).

### Publically available NIDM-Results packs

A set of 244 NIDM-Results packs has been made publically available on NeuroVault at http://neurovault.org/collections/1435/. Those packs describe the results of fMRI analyses performed at the subject (232 packs) and group (12 packs) levels on six datasets downloaded from OpenfMRI (RRID:SCR_005031) ^40,41^ (ds000005 1.1.0, ds000008 1.1.1, ds000011 unrevisioned, ds000052 unrevisioned, ds000107 unrevisioned, ds000114 unrevisioned).

### Examples of usage

#### Meta-analysis

From 21 pain studies (10 analysed in SPM and 11 in FSL) represented in NIDM-Results we performed group coordinate-based and image-based meta-analyses contrasting the effect of pain. The data and Python script used to perform these meta-analyses are available on NeuroVault (http://neurovault.org/collections/1425/ and GitHub ^42^ respectively.

Fig. 4 provides a schematic overview of the different steps involved to compute the meta-analyses. A set of NIDM-Results packs is queried in order to retrieve the information of interest that is then combined to perform the meta-analysis. Because the studies included in this meta-analysis are from a curated collection of pain studies from one laboratory, no manual filtering was needed for study or participant selection, and contrast selection was performed based on the contrast name.

**Fig. 4.**
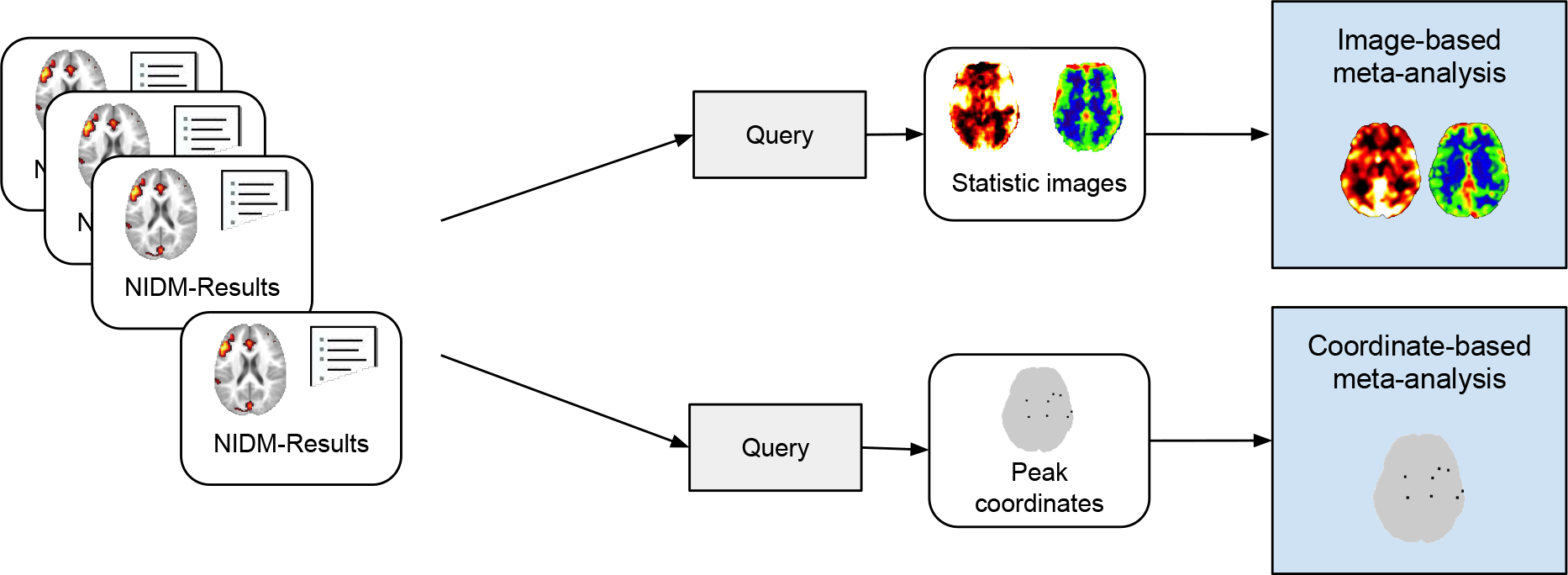
Image-based and coordinate-based meta-analyses using NIDM-Results. Each NIDM-Results pack is queried to retrieve the data and metadata of interest for each type of meta-analysis. These data are then combined in a meta-analysis.

The image-based meta-analysis was performed by combining the contrast estimate maps, along with their standard error, in a third-level mixed-effects general linear model (GLM). Each NIDM-Results pack was queried to retrieve the image data needed for the meta-analysis (i.e. the contrast image and contrast standard error image) along with the analysis mask. The query used to extract these data is displayed in Fig. 5. The name of the corresponding contrast was associated to each map to allow for the selection of the appropriate contrast. The neuroimaging software package used for the analysis was also extracted in order to identify which study estimates would need re-scaling. Second, the contrast and standard error estimates were selected according to the contrast name, re-scaled if needed and combined in a mixed-effects GLM. Areas of significant activation (p<0.05 FWE cluster-wise with a cluster-forming threshold of p<0.001 uncorrected) found by the pain meta-analysis are displayed in Fig. 6. Results are also available on NeuroVault at http://neurovault.org/collections/1432/.

**Fig. 5.**
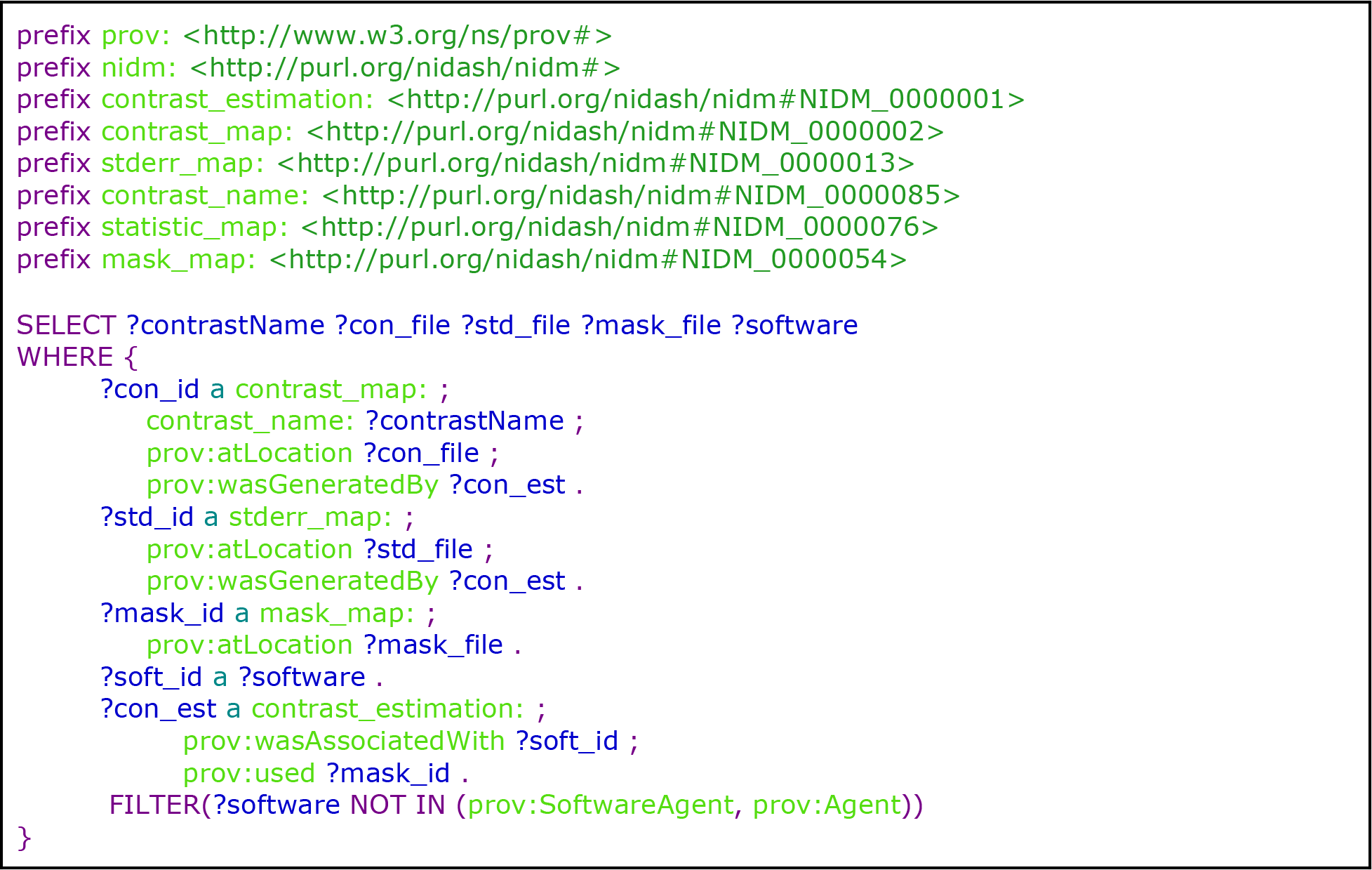
SPARQL query to retrieve data and metadata needed for image-based meta-analysis (syntax was highlighted using CodeMirror 66)

**Fig. 6.**
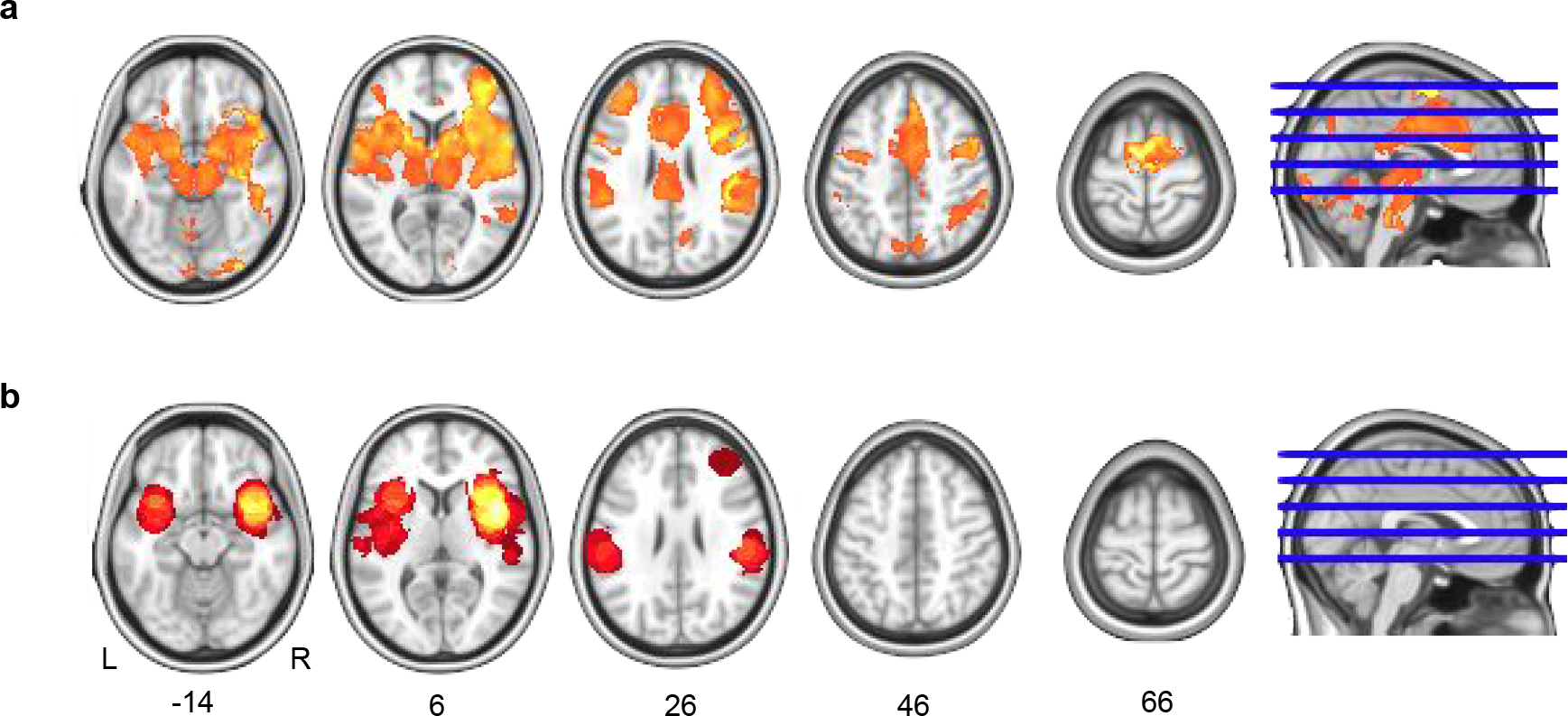
One-sample meta-analysis of 21 studies investigating the effect of pain. Areas of significant activation with an FWE-corrected cluster-wise threshold p<0.05 (cluster-forming threshold p<0.001 uncorrected) for the image-based (A) and the coordinate-based (B) meta-analyses.

The coordinate-based meta-analysis was performed using a Multilevel Kernel Density Analysis (MKDA) ^2^. Each NIDM-Results pack was queried to retrieve the coordinates of the local maxima, the reference space in use and the number of subjects for each contrast. Areas of significant activation (p<0.05 FWE cluster-wise with a cluster-forming threshold of p<0.001 uncorrected) found by the pain meta-analysis are displayed in Fig. 6.

In line with previous results from the literature ^4^, the detections for the coordinate-based and image-based meta-analysis appear consistent with a lower sensibility of the coordinate-based meta-analysis.

#### Reporting of neuroimaging results

Table 2 provides a mapping between the guidelines provided in ^8^ to report neuroimaging results and the fields available in a NIDM-Results serialisation. NIDM-Results cover all elements from the “Statistical modelling” checklist that could be automatically retrieved within the neuroimaging software package.

**Table 2.**
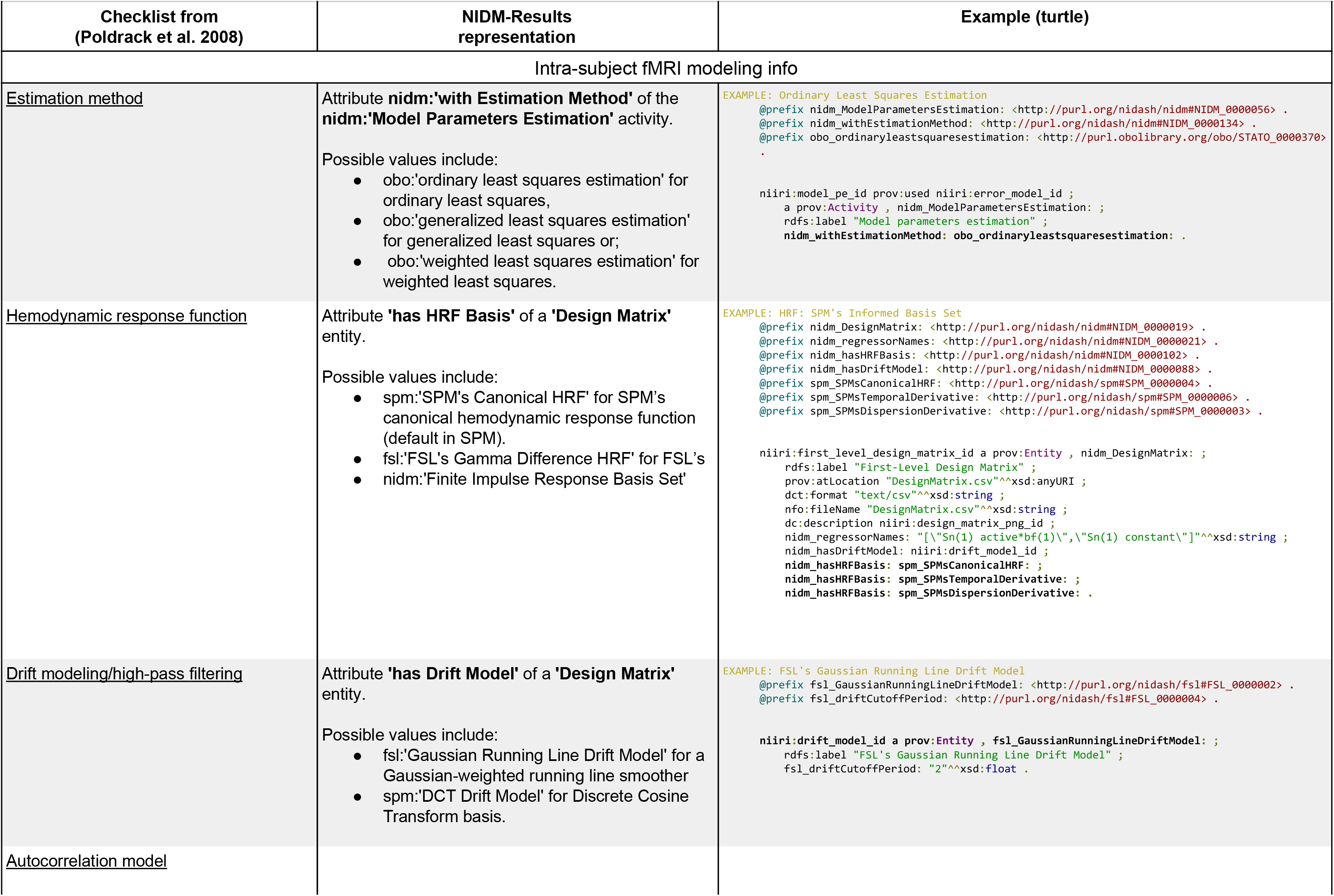
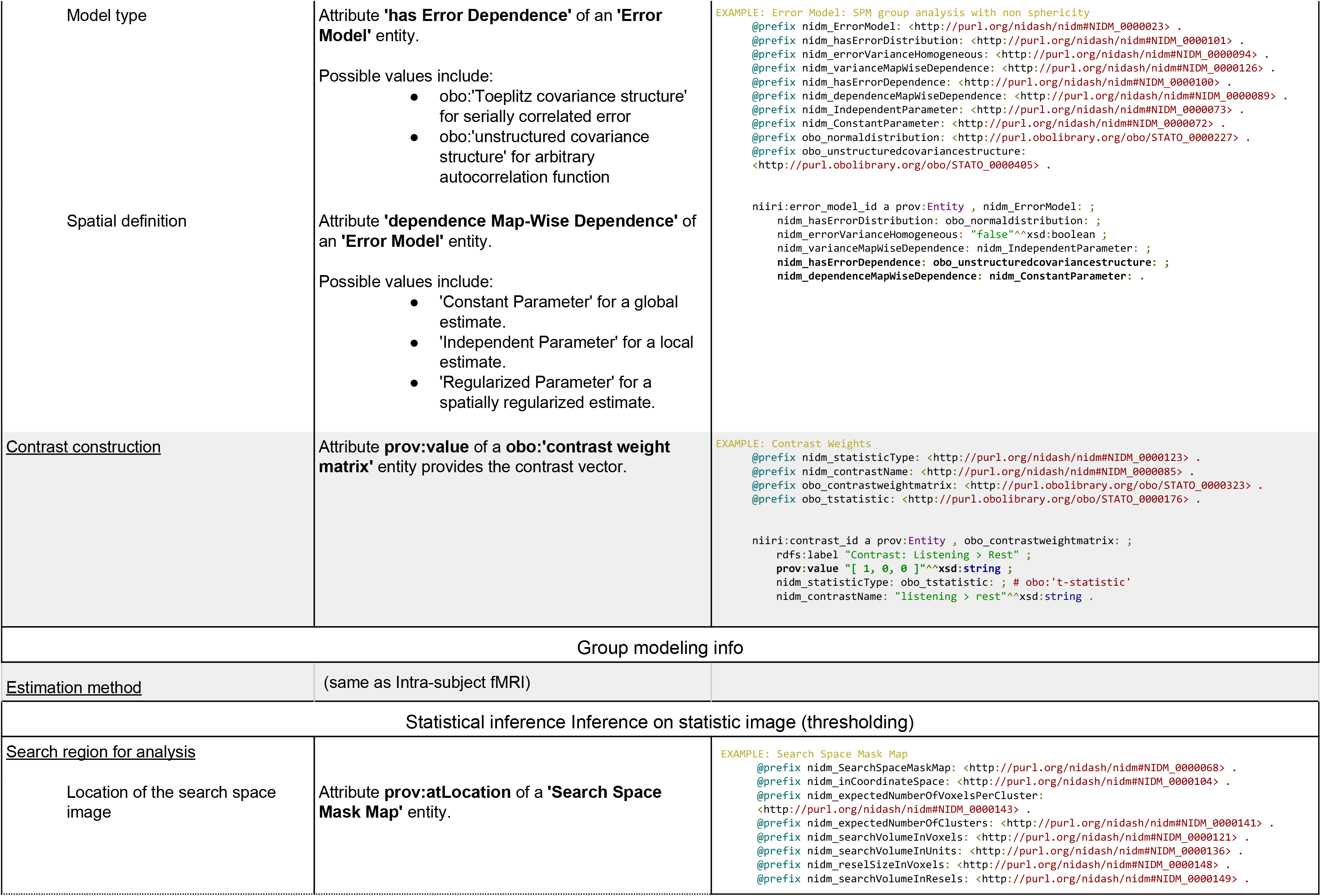
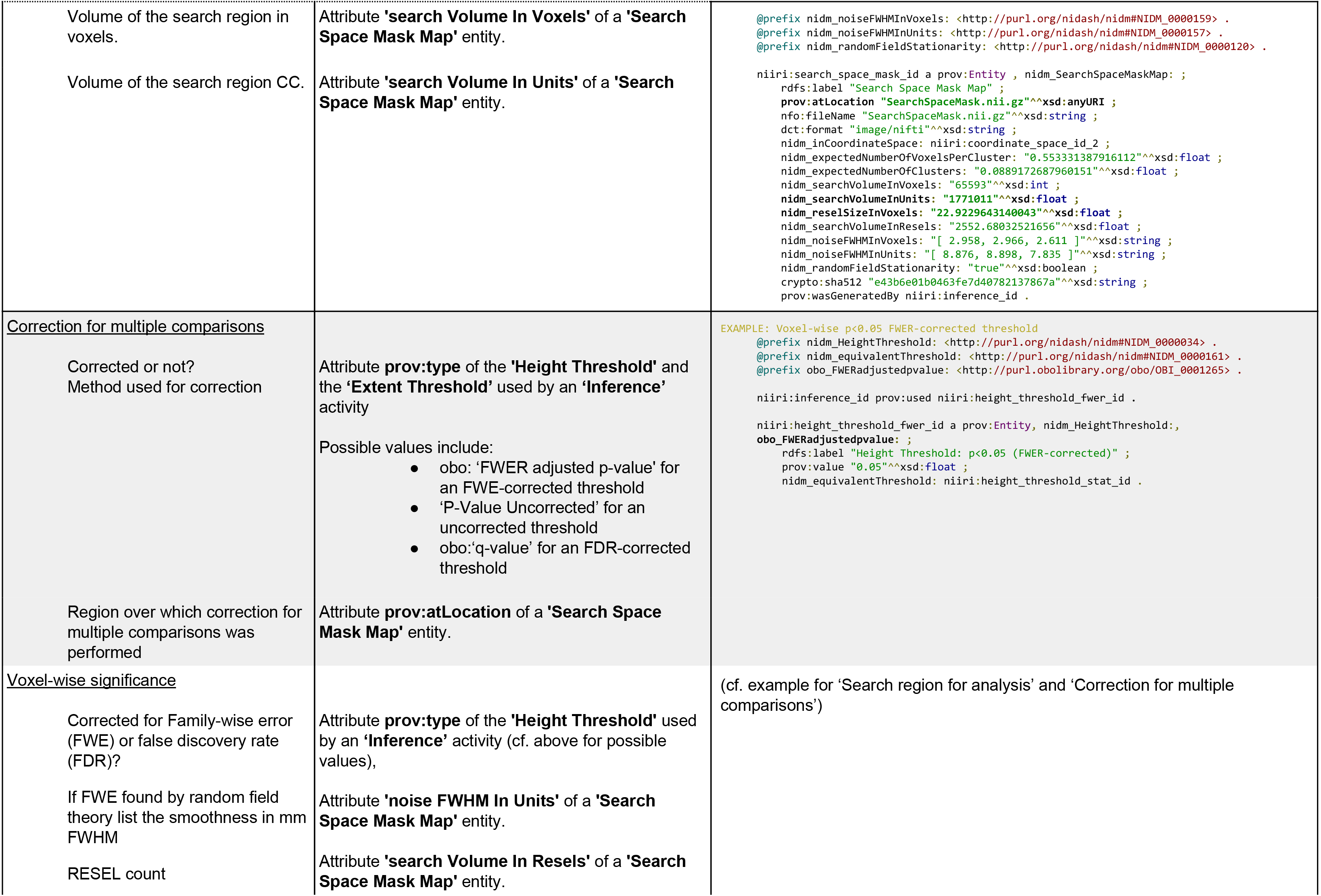
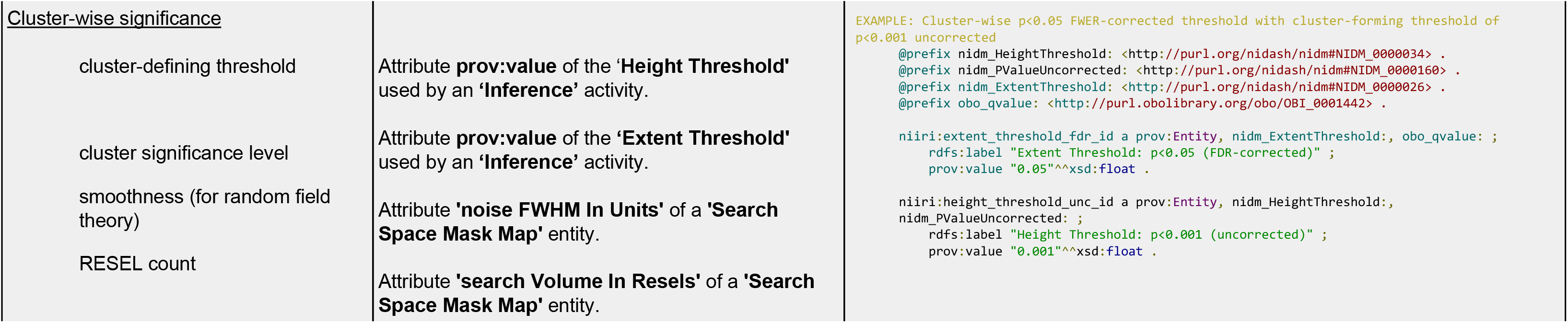
Checklist to report neuroimaging results for intra-subject fMRI and group models from (Poldrack et al. 2008) and corresponding representation in NIDM-Results. The following items from the original checklist were excluded as not available automatically: “design type”, “orthogonalization of regressors”, “additional regressors used”,”if not whole brain, state how region of analysis was determined”, “If correction is limited to a small volume, the method for selecting the region should be stated explicitly”, “threshold used for visualization in figures”, “correction for multiple planned comparisons within each voxel”.

**Table 3.** Prefixes of the vocabularies used in NIDM-Results.

| Vocabulary/Ontology | URI | Prefix |
| --- | --- | --- |
| PROV | http://www.w3.org/ns/prov# | prov |
| STATO | http://purl.obolibrary.org/obo/ | obo |
| NeuroLex | http://uri.neuinfo.org/nif/nifstd/ | nlx |
| RRID | http://scicrunch.org/resolver/ | rrid |
| Dublin Core types | http://purl.org/dc/dcmitype/ | dctype |
| Dublin Core elements | http://purl.org/dc/elements/1.1/ | dc |
| Dublin Core terms | http://purl.org/dc/terms/ | dct |
| Cryptographic Hash Functions | http://id.loc.gov/vocabulary/preservation/cryptographicHashFunctions# | crypto |
| NEPOMUK file ontology | http://www.semanticdesktop.org/ontologies/2007/03/22/nfo# | nfo |
| NIDM | http://purl.org/nidash/nidm# | nidm |
| FSL | http://purl.org/nidash/fsl# | fsl |
| SPM | http://purl.org/nidash/spm# | spm |

Examples of reports generated from a NIDM-Results export of group and single-subject analyses performed in SPM and FSL are provided in Fig. 7. The data and Python script used to generate those report are available on NeuroVault (http://neurovault.org/collections/1435/) and GitHub ^42^ respectively.

**Fig. 7.**
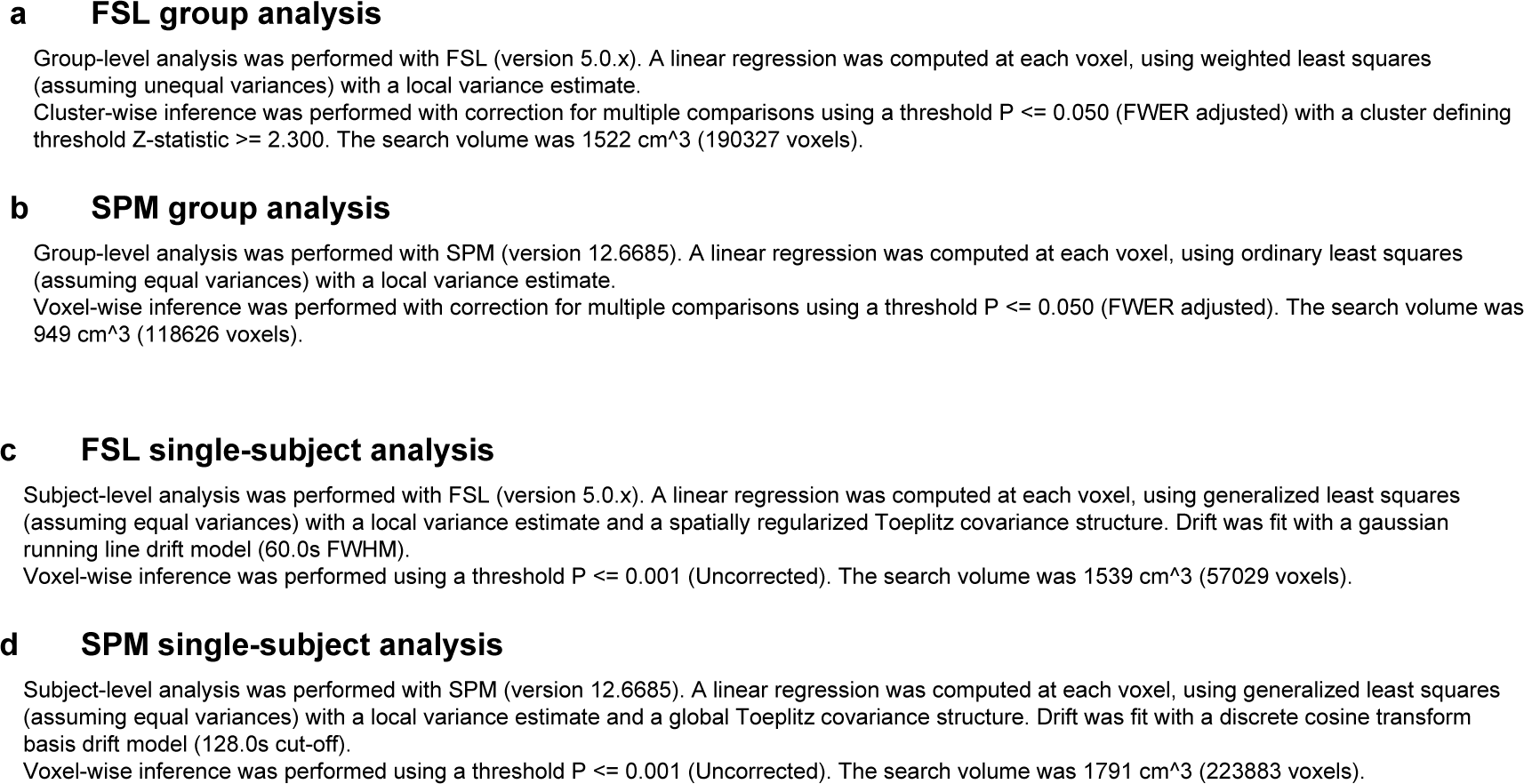
Examples of reports generated from NIDM-Results packs for group (a, b) and single-subject (c, d) analyses performed in FSL (a, c) and SPM (b, d).

## Discussion

Data sharing in the neuroimaging community is still restrained by a number of psychological and ethical factors that are beyond the scope of the current paper (see ^7^,43 for a review). Those will have to be addressed in order for data sharing to become common practice in the neuroimaging community. In an effort to address the technological barriers that make data sharing challenging, here we have proposed a solution to share neuroimaging results of mass univariate analyses.

As a first step to provide machine-readable metadata, we restricted our scope to information that was automatically extractable and attributes that were crucial for meta-analysis (e.g. number of subjects). This limited the amount of information that could be represented. For instance, the description of the paradigm was limited to the design matrix and a list of regressor names. Ideally, to be able to automatically query for studies of interest, one would need a more thorough description of the paradigm and of the cognitive constructs involved. While vocabularies are becoming available (e.g. Cognitive Atlas ^44,45^ and CogPO ^46,47^), description of fMRI paradigms is still a topic of active research. Some level of manual interaction to select contrasts of interest is therefore needed to compute a meta-analysis based on NIDM-Results packs. Nevertheless, NIDM-Results allows for the automation of part of the meta-analysis as described in our results. In the future, as a consensus develops on the description of paradigms, NIDM-Results could easily be extended to include this information. Similarly, NIDM-Results could be extended to match emerging best practices (such as ^10^).

NIDM-Results currently focuses on the representation of parametric mass-univariate analyses. Thanks to the intrinsic extensibility of RDF models, variants could be proposed to broaden its scope. For example, an extension for non-parametric statistics is under discussion ^48^. Mass-univariate results, as the most well established approach for fMRI analysis, was an obvious choice to start a cross-software modelling effort. But neuroimaging cannot be limited to mass-univariate analyses and future work will focus on providing extensions for other types of analyses (e.g. analysis of resting state fMRI).

We based our modelling effort on PROV, a specification endorsed by the W3C, to model provenance on the web. Other efforts have been proposed to model provenance including families of ontologies like the OBO foundry ^49^ or DOLCE ^50^. We chose PROV as it is lightweight, focused only on provenance, and is easily extensible to provide domain-specific knowledge.

Another recent effort to provide structured organisation of neuroimaging data is the Brain Imaging Data Structure (BIDS)^51^. While NIDM-Results and BIDS both concern the organisation and description of neuroimaging data, they operate at very distinct domains of the analysis path. BIDS provides a mechanism for organising only the original raw data, and it does not cover any derived data nor the definition of particular statistical models or the outputs of those models. NIDM-Results, in contrast, works at the other end of the analysis pipeline, defining a framework to describe the statistical model, the statistical ‘contrasts’ that interrogate the model, and the resulting statistical maps and inferences obtained from each contrast. Whereas BIDS was designed so that an end-user could manually create the files and directories of a BIDS-compliant data structure, NIDM--Results is intended to be automatically generated from analysis software and was therefore created using more expressive semantic web technologies. Of the larger NIDM project, it is the NIDM-Experiment portion that will have the greatest overlap with BIDS. By making the experimental metadata available as linked data, NIDM-Experiment will enable querying across the full neuroimaging data lifecycle, interrogating data possibly hosted on distributed resources.

NIDM-Results is based on RDF and semantic web technologies. While a number of ontologies have been developed in relation with neuroimaging (e.g. Cognitive Atlas ^44^, CogPO ^46^, OntoNeurolog ^52^), the use of controlled vocabularies and of linked data is not yet common practice in our community. As more and more data become available online and as standardisation effort like the RII develops, we believe that these technologies will become more widespread. RDF was chosen as a basis for NIDM for the expressivity of its graph-based structures, the possibility to form intricate queries across datasets ^37^, as well as for the extensibility of the created data models and the possibility to interconnect across knowledge domains (cf. ^53^ for a review).

One limitation of NIDM-Results is that only limited provenance is represented. For instance, computational environment, which has been shown to be source of undesired variability in neuroimaging results ^54^, is not part of our model. NIDM-Results is part of a broader effort (NIDM) that aims at representing different levels of provenance in neuroimaging experiments. While those efforts are still under development, our goal is to keep a link between those components to eventually provide a complete representation of neuroimaging provenance.

As for the definition of any new model, gaining acceptance within the neuroimaging community will be crucial for NIDM-Results. To insure a level of consensus, including the point-of-view of different actors in the field, NIDM-Results was built as part of a collaborative effort. More feedback from the community is welcome and can be submitted as issues in our GitHub repository or by email at. We also made a strong commitment to make implementations available. Taking advantage of the fact that most functional MRI studies are performed using a limited number of software packages (> 75% for SPM or FSL, > 90% for SPM, FSL or AFNI according to ^55^), we developed implementations for SPM and FSL, and are currently working with AFNI developers to further extend the coverage of NIDM-Results export.

While we have focused our implementation efforts on the generation of NIDM-Results packs, the development of applications processing NIDM-Results is also crucial, to serve as incentives for neuroimaging users. As an example, we liaised with NeuroVault to propose a one-click upload of NIDM-Results. Here, users can benefit from all NeuroVault features including state-of-the-art visualisations but also sharing, either privately or publicly depending on the stage of the project. This process can ease communication between researchers working on different platforms or used to a different set of neuroimaging tools. In the future, we plan to offer a one-click upload of NIDM-Results packs from the neuroimaging software packages (SPM, FSL, AFNI) into NeuroVault. A wider ecosystem is also under development (including a standalone viewer).

Future work will focus on developing extensions for NIDM-Results to cover a larger spectrum of neuroimaging studies (e.g. non-parametric analyses) as well as to stay up-to-date with emerging best practices. We will also sustain our effort on the development of tools that can read and write NIDM-Results. Finally we will focus on the specification of the other NIDM components to enable modelling of a complete fMRI experiment from raw data to statistical results.

We believe NIDM-Results is an essential tool for the future of transparent, reproducible science using neuroimaging. If all research publications were accompanied by such a machine-readable description of the experiment, debates on the exact methodology used would be compressed or eliminated, and any replication efforts greatly facilitated.

## Methods

### Process

Since August 2013 the model was developed through weekly teleconferences and eight focused workshops during which the team of experts iteratively defined the terminology, seeking to ensure that the output of AFNI, FSL and SPM could be represented in this framework. Furthermore, a separate meeting was organised with each of the development teams of SPM, FSL and AFNI to discuss the model and its implementation. Minutes of the meetings and online discussions are publicly available in our shared Google drive ^56^ and on GitHub under the incf-nidash organization ^57^.

### Scope of the model

NIDM-Results focuses on mass-univariate models based on a General Linear Model (at the subject or group level). To facilitate adoption, we restricted the scope of NIDM-Results to metadata that could be automatically extracted with limited user input, motivated by the specific metadata that is crucial for the application of meta-analysis. This had important practical consequences. Given that pre-processing and statistical analysis are sometimes done using separate pipelines, we focused on the statistical analysis only. The concepts to be represented in NIDM-Results were selected based on (1) meta-analysis best practices; (2) published guidelines to report fMRI studies ^8^, and (3), in an effort to ensure continuity with current practice, we also considered the elements displayed as part of results reporting in different neuroimaging software (e.g. peaks, clusters). When an item, essential for image-based meta-analysis, was not produced as part of the standard analysis (e.g. the contrast standard error map in SPM) we included it in the model and depend on the exporters to generate it from existing data.

### Term re-use and definitions

For each piece of information, we checked if an appropriate term was available in publicly available ontologies: in particular STATO for statistics term, PROV for provenance, NeuroLex for neuroscience terms, RRID for tools and also, to a lesser extent, Dublin Core, the NEPOMUK file ontology and the Cryptographic Hash Functions vocabulary. Namespaces of the re-used ontologies are provided in Table 2. More details on the re-used vocabularies are provided below.

#### PROV

The W3C specification PROV ^35^ defines three types of objects: an *Activity* represents a process that was performed on some data (e.g. a voxel-wise inference) and occurred over a fixed period of time; an *Agent* represents someone (human, organization, machine…) that takes responsibility for an activity (e.g. the SPM software) and, finally, an *Entity* represents any sort of data, parameters etc. that can be input or output of an activity (e.g. a NIfTI image). PROV also defines a set of relations between those objects (e.g. a voxel-wise inference *Activity used* a NIfTI image *Entity*; a voxel-wise inference *Activity was associated with* the SPM *Agent* and another NIfTI image *Entity was generated by* the segmentation *Activity*). NIDM-Results terms were defined as specialisations of PROV terms.

#### STATO

GLM analyses of fMRI data rely on well-known statistical constructs (e.g. one-sample T-test, two-sample T-test, F-tests, ANOVA, inference, ordinary least squares estimation, etc.). The general-purpose STATistics Ontology (STATO) ^58^ is built on the top of the OBO foundry and aims to provide a set of terms describing statistics. We re-used statistics terms available in STATO (e.g. obo:’t-statistic’) and when we could not find an appropriate statistical term, we engaged with STATO developers through GitHub issues to propose new terms (e.g. “residual mean squares” discussed in issue 35 ^59^).

#### NeuroLex and RRID

Much work has been done in the neuroimaging community to provide controlled vocabularies and ontologies defining neuroimaging concepts. NeuroLex ^60,61^ provides a common platform that gathers terms from different sources (including previous vocabularies developed by NIF, BIRN…). Interestingly, Neurolex was part of the recent Resource Identification Initiative (RII) ^62,63^ that publicized the use of those identifiers (e.g. “RRID:SCR_007037” for SPM ^64^) in research papers. RII is currently focused on the identification of biological resources and has been quickly adopted, with more than 100 journals participating to date. We re-used the available RRIDs describing neuroimaging software packages.

#### Dublin Core, NEPOMUK file ontology and the Cryptographic Hash Functions vocabulary

Many vocabularies and ontologies have terms available to describe files. We chose to rely on the widely adopted DUBLIN core terminology ^65^. Additionally, we used the “fileName” term from the NEPOMUK file ontology ^66^ and the SHA-256 term from the Cryptographic Hash Functions vocabulary ^67^.

#### New terms

When no term was found to describe a given neuroimaging concept of interest, we created a new term and carefully crafted a definition or engaged with the relevant ontology maintainers (e.g. we contributed 41 terms to STATO) to propose a new definition. All new terms and definitions were thoroughly discussed between our panel of experts in the NIDM working group, which is part of the INCF Neuroimaging Task Force (NIDASH).

### Examples of usage

#### Meta-analysis

Data collection was subject to the Oxford University ethics review boards, who approved the experiments. Only statistical summaries with no identifying data are shared along this manuscript.

Results from 21 pain studies previously analysed with FSL were made available to us. The second-level analyses were recomputed with SPM for 10 of those studies in order to obtain a dataset of NIDM-Results packs coming from mixed software packages. We computed a one-sample meta-analysis contrasting the effect of “pain” and compared the results of coordinate-based and image-based meta-analyses.

The MKDA toolbox ^68^, was used to perform the coordinate-based meta-analysis. The nidmresults Python toolbox (https://pypi.python.org/pypi/nidmresults) was used to generate the csv file required as input for the analyses.

FSL’s FLAME 1 ^69^ was used to compute the image-based meta-analysis with the gold standard approach (3rd level mixed-effects general linear model). FLAME 1 implements a random-effects meta-analysis with iterative estimation of between-study variation via maximum likelihood ^70,71^. Parametric inferences are conducted by reference to a Student’s t distribution with nominal degrees of freedom (i.e. number of studies minus number of regression parameters) to account for uncertainty in the estimation of the between-study variance parameter. Difference in data scaling between software packages were compensated by rescaling the FSL maps to a target intensity of 100 (instead of 10 000 by default).

#### Reporting of neuroimaging results

From four studies exported with NIDM-Results we wrote a script ^42^ to extract the information of interest to describe group and subject-level statistics using the RDFlib library ^72^ to query the documents. The paragraph that was generated could, for instance, be used as part of the method section in a research paper.

## Acknowledgments

We gratefully acknowledge Matthew Webster, Paul McCarthy, Eugene Duff and Steve Smith, from the FMRIB and Robert Cox and Ziad Saad from the NIH, for their inputs on the integration of NIDM-Results within FSL and AFNI; as well as the NIDASH task force members for their inputs during the development of the model. We also gratefully acknowledge the Tracey group at FMRIB for sharing their pain datasets used in the meta-analysis.

## Author Contributions

CM contributed in creation of NIDM-Results, developed the FSL exporter and wrote the manuscript.

TA contributed in creation of NIDM-Results and edited the manuscript.

AB generated the example datasets and edited the manuscript.

GC provided feedback on the implementation of the model for AFNI and commented on the manuscript.

SD contributed in creation of NIDM-Results and edited the manuscript.

GF contributed in creation of NIDM-Results, developed the SPM exporter and edited the manuscript. SG contributed in creation of NIDM-Results and edited the manuscript.

TG contributed in creation of NIDM-Results, developed the CBRAIN plugin and edited the manuscript.

KJG contributed in creation of NIDM-Results, integrated NIDM-Results with NeuroVault and edited the manuscript.

KGH contributed in creation of NIDM-Results and edited the manuscript.

MJ provided feedback on the implementation of the model for FSL and commented on the manuscript.

DBK contributed in creation of NIDM-Results and edited the manuscript.

BNN contributed in creation of NIDM-Results and edited the manuscript.

JBP contributed in creation of NIDM-Results and edited the manuscript.

RR provided feedback on the implementation of the model for AFNI and commented on the manuscript.

VS developed a viewer for NIDM-Results and edited the manuscript.

JT contributed in creation of NIDM-Results and commented on the manuscript.

TEN contributed in creation of NIDM-Results and edited the manuscript.

